# A network algorithm for the X chromosomal exact test for Hardy-Weinberg equilibrium with multiple alleles

**DOI:** 10.1101/2020.09.20.305102

**Authors:** Jan Graffelman, Leonardo Ortoleva

## Abstract

Statistical methodology for testing Hardy-Weinberg equilibrium at X chromosomal variants has recently experienced considerable development. Up to a few years ago, testing X chromosomal variants for equilibrium was basically done by applying autosomal test procedures to females only. At present, male alleles can be taken into account in asymptotic and exact test procedures for both the bi- and multiallelic case. However, current X chromosomal exact procedures for multiple alleles rely on a classical full enumeration algorithm and are computationally expensive, and in practice not feasible for more than three alleles. In this article we extend the autosomal network algorithm for exact Hardy-Weinberg testing with multiple alleles to the X chromosome, achieving considerable reduction in computation times for multiallelic variants with up to five alleles. The performance of the X-chromosomal network algorithm is assessed in a simulation study. Beyond four alleles, a permutation test is, in general, the more feasible approach. A detailed description of the algorithm is given and examples of X chromosomal indels and microsatellites are discussed.

## 1 Introduction

The statistical testing of genetic variants for Hardy-Weinberg equilibrium (HWE) is an important part of the analysis of genetic datasets, for a variety of reasons. Gross deviations from equilibrium are often the result of genotyping errors, and testing can be helpful to detect such errors (Hosking et al., 2004; Teo et al., 2007; Leal, 2005; Chen et al., 2017). Moreover, many methods used in genetic data analysis rely on the equilibrium assumption, and the filtering of variants on the basis of their p-values obtained in test for HWE can be used as a safeguard to prevent violation of assumptions made. A recent overview of statistical tests for Hardy-Weinberg equilibrium is given by Graffelman (2020). Currently, exact test procedures are the state-of-the-art for testing biallelic genetic variants, and are most commonly employed. Fast recursive procedures are available that can do exact testing of biallelic variants for HWE on a genome-wide scale (Wigginton et al., 2005; Chang et al., 2015). For variants with multiple alleles the exact test is computationally more costly. Algorithms for the efficient exact testing of multiallelic variants have been proposed by several authors (Louis and Dempster, 1987; Guo and Thompson, 1992; Huber et al., 2006). A recursive network algorithm (Aoki, 2003; Engels, 2009) has been proposed for more efficient calculation of exact p-values. When the computational cost of the network approach becomes prohibitive, a permutation test based on the sampling of outcomes from the exact distribution can still be used as an alternative in order to obtain an approximate p-value (Guo and Thompson, 1992).

Recently, the statistical testing for HWE of variants on the X chromosome has experienced considerable development. Up to a few years ago, X chromosomal variants were tested using autosomal procedures for the females only. Graffelman and Weir (2016) developed the full suit of frequentist test procedures (a two degrees of freedom asymptotic chi-square test, an exact test and a permutation test) specifically for X chromosomal variants, which take male alleles on X into account. This has the advantage of a larger sample size (more X chromosomes), implying higher precision for the estimation of allele frequencies, and the potential rejection of equilibrium when there is a difference in allele frequency between the sexes. The biallelic exact test for X is computationally feasible for a complete X chromosome, and efficient C++ code for this test is currently shared by the PLINK software (Purcell et al., 2007; Chang et al., 2015) and the R-package HardyWeinberg (Graffelman, 2015). Later, Graffelman and Weir (2018) extended their X chromosomal exact test for multiple alleles with a classical full enumeration algorithm, and reported on the analysis of all triallelic variants on X at considerable computational cost, and suggesting the use permutation tests based on sampling for X chromosomal variants with four or more alleles. In this article, we propose a modification of the network algorithms proposed by Aoki (2003) and Engels (2009), adapting the network algorithm for the X chromosome. The network algorithm efficiently avoids the repeated calculation of factorial terms that are shared in the list of possible outcomes generated by complete enumeration, leading to large computational savings. This way, we strive to extend the application of the X chromosomal exact test towards variants with a larger number of alleles while maintaining computation time within feasible limits.

The structure of the remainder of this article is as follows. In Section 2 we review exact tests with multiple alleles for the autosomes and for the X chromosome, and present the adaptation of the network algorithm to the X chromosome. In Section 3 we assess the performance of the new network algorithm in a simulation study. Section 4 shows examples of the network based test for a varying number of alleles with data taken from the 1,000 genomes project (The 1000 Genomes Project Consortium, 2015). We describe the HWE analysis of a complete X chromosome of the Tuscan population (TSI) of the 1,000 genomes project, and also address the analysis of a forensic database of X-chromosomal microsatellites (Chen et al., 2018). The Discussion in Section 5 completes the manuscript.

## 2 Theory

In this section we review exact inference for HWE with multiple alleles, and explain the operation of the network algorithm for both the autosomal and X chromosomal case with a toy example.

### 2.1 Autosomal exact inference with multiple alleles

Exact inference for autosomal variants with multiple alleles is based on the conditional distribution of the genotype counts, considering all observed allele counts as given. This distribution was derived by Levene (1949), and is given by

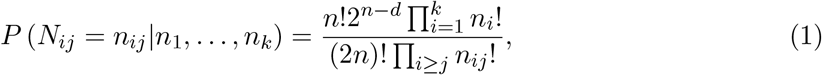

where *n* represents the sample size, *n_i_* the count of the *i*th allele, the count of genotype *ij*, and *d* = ∑*n_ii_* the total homozygote frequency. Eq. 1 describes the distribution of the genotype counts under the assumption of HWE. One first calculates the probability of the observed sample according to Eq. 1. Next, a full enumeration is made of all possible genotype arrays that are compatible with the observed total allele counts, and their probabilities are calculated. Finally, the exact p-value is obtained by summing the probabilities of all genotype arrays that are less likely than or equally likely to the observed sample. The full enumeration approach combined with the calculation of Eq. 1 is computationally expensive, in particular for large samples with many alleles. A full enumeration algorithm for an arbitrary number of alleles has been described by Louis and Dempster (1987).

A drawback of the classical full enumeration algorithm is that many genotype arrays involve the same factorials, which will be repeatedly calculated if a simple loop is used to iterate over all possible arrays. The network algorithm enables the sharing of the calculation of common factorials across similar genotype arrays, and can so produce considerable computational savings. The network approach was proposed by Mehta and Patel (1983) who developed this algorithm for a more efficient calculation of Fisher’s exact test for large contingency tables. Aoki (2003) presented the first network algorithm for exact testing in the context of Hardy-Weinberg equilibrium. The computation of Eq. 1 is simplified by recognising that for given allele counts, the part

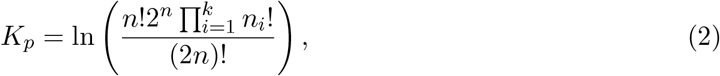

is a common factor for all genotype arrays, and can taken as a constant which is calculated only once. To further simplify the calculations, we take the logarithm of Eq. 1 and have

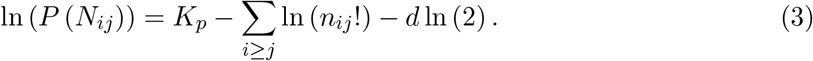

Figure 1 shows the operation of an autosomal network algorithm for a triallelic variant, with a sample of size *n* = 8 and allele counts 7, 5 and 4 for A, B and C respectively. Each path from left to right generates a particular genotype array. The probability of the sample generated by the traced path is 0.028, and if this sample is observed, the exact test p-value is 0.167. The network has 21 paths corresponding to 21 different possible genotype arrays for the given allele counts.

**Figure 1:**
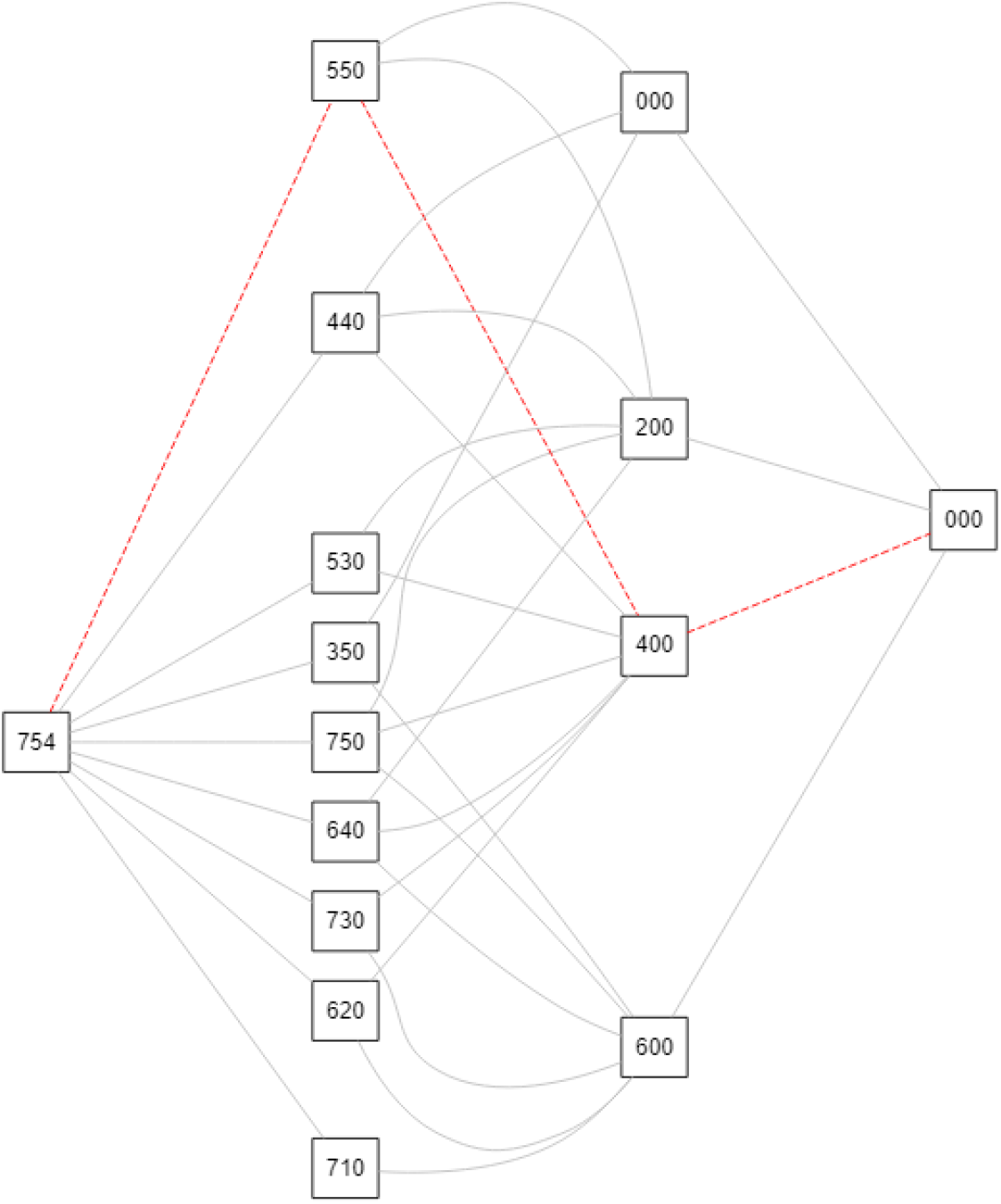
Graph for an autosomal triallelic variant for a sample size of *n* = 8 with allele counts (A=7,B=5,C=4). Nodes represent allele counts, and edges the assignment of alleles to genotypes. The dashed path illustrates the generation of 1 CC homozygote and 2 AC heterozygotes, leaving allele counts (A=5,B=5,C=0), followed by the generation of two BB homozygotes and one AB heterozygote, leaving (A=4,B=0,C=0), and finally the generation of two AA homozygotes to arrive at (A=0,B=0,C=0). The generated genotype array is (AA=2,BB=2,CC=1,AB=1,AC=2,BC=0). Each path in the network traces the generation of a genotype array that is compatible with the observed allele counts. The network exhausts all possible genotype arrays for the given allele counts.

The algorithm proceeds by computing the second term of logfactorials in Eq. 3 incrementally, exhausting alleles one by one. As Fig. 1 shows, the edge from 754 to 550 is shared by three genotype arrays, and the corresponding logfactorials of *n_AA_* and *n_AC_* only need to be computed once. For more details on the autosomal network algorithm we refer to Aoki (2003) and Engels (2009).

### 2.2 X chromosomal exact inference with multiple alleles

For exact testing for Hardy-Weinberg equilibrium at X chromosomal variants with multiple alleles, Graffelman and Weir (2018) derived, assuming equality of allele frequencies in the sexes and Hardy-Weinberg proportions in females, the exact distribution of the joint distribution of the number of female heterozygotes and the number of hemizygous males given by

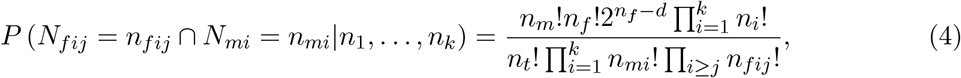

where *n_m_* and *n_f_* represent the numbers of males and females, *n_t_* = 2*n_f_* + *n_m_* the total number of alleles, *n_mi_* and *n_fij_* male and female genotype counts and *d* the total number of homozygote females. To show the increase in computational complexity, we use the same set of allele counts (A=7,B=5,C=4) as in Fig. 1, but now consider gender, assuming the sample is composed of 4 males and 6 females, totalling 16 alleles. Figure 2 shows the network for this case, and the construction of the genotype array (*m_A_* = 3, *m_B_* = 1, *m_C_* = 0, *f_AA_* = 1, *f_BB_* = 2, *f_CC_* = 1, *f_AB_* = 0, *f_AC_* = 2, *f_BC_* = 0) is indicated. The number of possible genotype arrays, 136, has increased considerably in comparison with the previous autosomal variant with the same total allele counts. We follow the same approach as before, now defining two constants *K_p_* and *K_m_* as

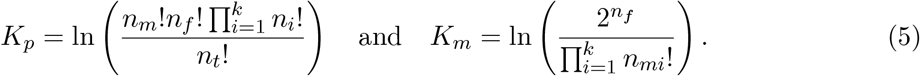

**Figure 2:**
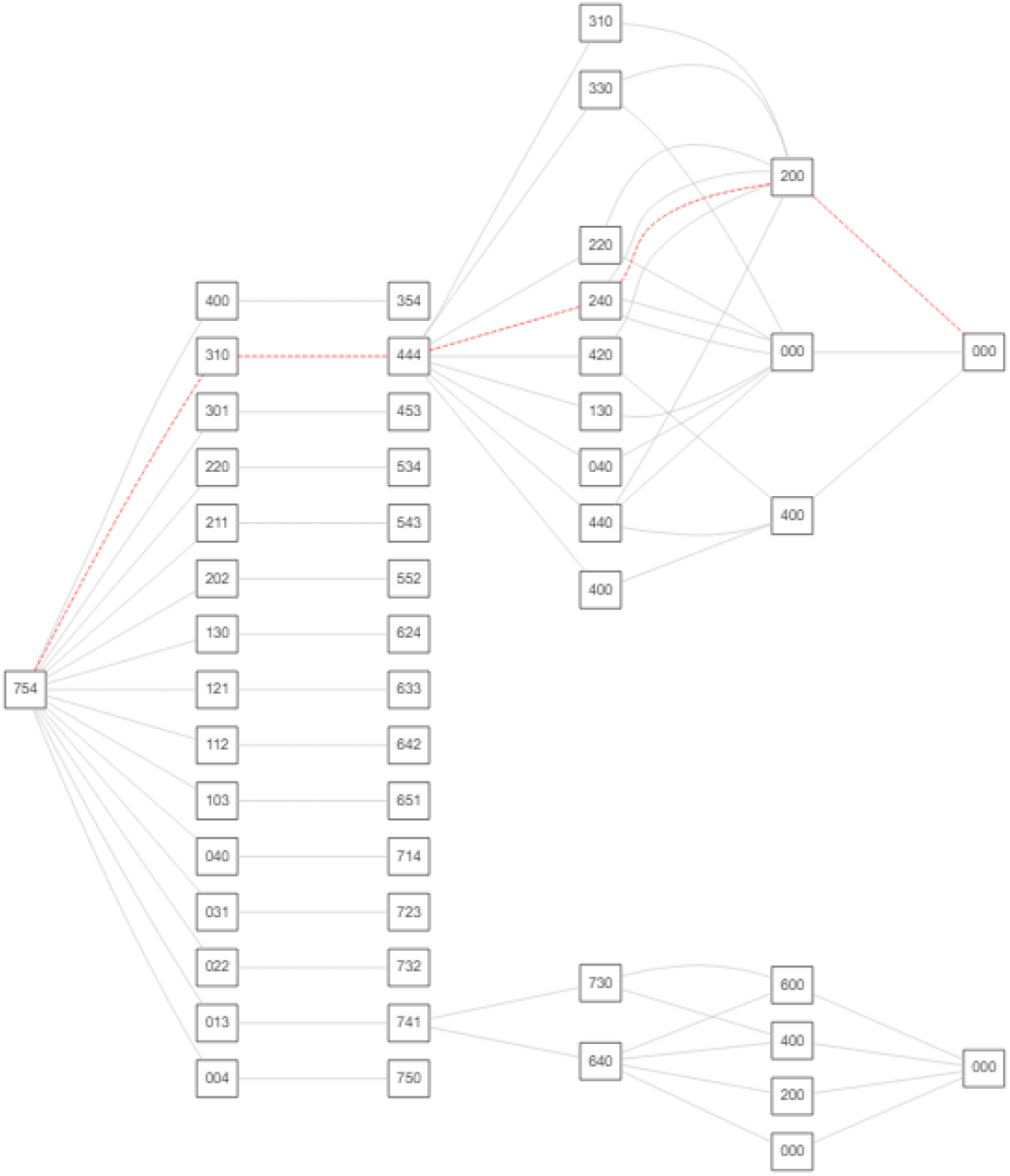
Network for an X chromosomal triallelic variant for a sample size of 4 males and 6 females with total allele counts (A=7,B=5,C=4). Nodes represent allele counts, and edges the assignation of alleles to genotypes. The dashed path illustrates the generation of 3 A males and 1 B male, followed by the generation of the females. The generated genotype array is (*m_A_* = 3, *m_B_* = 1, *m_C_* = 0, *f_AA_* = 1, *f_BB_* = 2, *f_CC_* = 1, *f_AB_* = 0, *f_AC_* = 2, *f_BC_* = 0). Each path in the network traces the generation of a genotype array compatible with the observed allele counts. The network exhausts all possible genotype arrays for the given allele counts and the given number of males and females. For simplicity the network for female genotypes is shown only for two sets of female allele counts.

Taking logarithms, one has

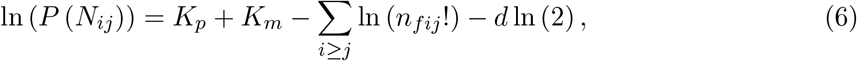

and again the sum of the logfactorials is incrementally evaluated one allele at a time. In essence, for X chromosomal variants we first generate all possible male genotype arrays, and next apply the autosomal network algorithm using the remaining female allele counts.

## 3 Simulation study

The X chromosomal exact test involves the computation of the probabilities of all possible genotype arrays for the given allele counts, according to Eq. 4. The number of arrays is in general, larger than for the autosomes, and the X chromosomal exact test is computationally expensive for systems with multiple alleles. We compare the computational cost of the network algorithm with a classical enumeration algorithm for bi- and triallelic variants, and also compare the network algorithm with a permutation test for three or more alleles. We expect the network algorithm to be computationally cheaper because it is able to store partial results thanks to recursion through the network, which avoids repeating calculations from the beginning for every possible table of genotypes under analysis, as explained in the previous section. Full enumeration algorithms for X chromosomal exact test are currently only available for bi- and triallelic variants. Figure 3A shows the computation time as a function of the number of biallelic X chromosomal SNPs. X chromosomal SNPs were simulated under the assumptions of Hardy-Weinberg proportions in females and equality of male and female allele frequencies, using a sample size of *n* = 100 consisting of 50 males and 50 females. All computations were carried out in the R environment (R Core Team, 2014), using a server with thirty-two compute nodes, half of the nodes were 16-core Intel Xeon E5-2630 systems (2.40GHz; 128 Gb RAM); the other half were 24-core Intel Xeon Gold 5118 (2.30GHz; 384Gb RAM). For two alleles, the network algorithm is seen to take more time in comparison with an enumeration algorithm. The computation time of the network algorithm is seen to increase, as expected, linearly with the number of SNPs, though most conveniently shown in a logarithmic scale as in Figs. 3A and 3B. We simulated up to 4M biallelic SNPs, because the 1,000 genomes project (The 1000 Genomes Project Consortium, 2015) reports about 3.5M biallelic SNPs on X. For biallelic SNPs, we used `HWExactStats` implementation of the enumeration algorithm, and the HWNetwork implementation of the network algorithm. The first uses C code shared with PLINK, and the latter modified C code of Engels’s autosomal algorithm.

**Figure 3:**
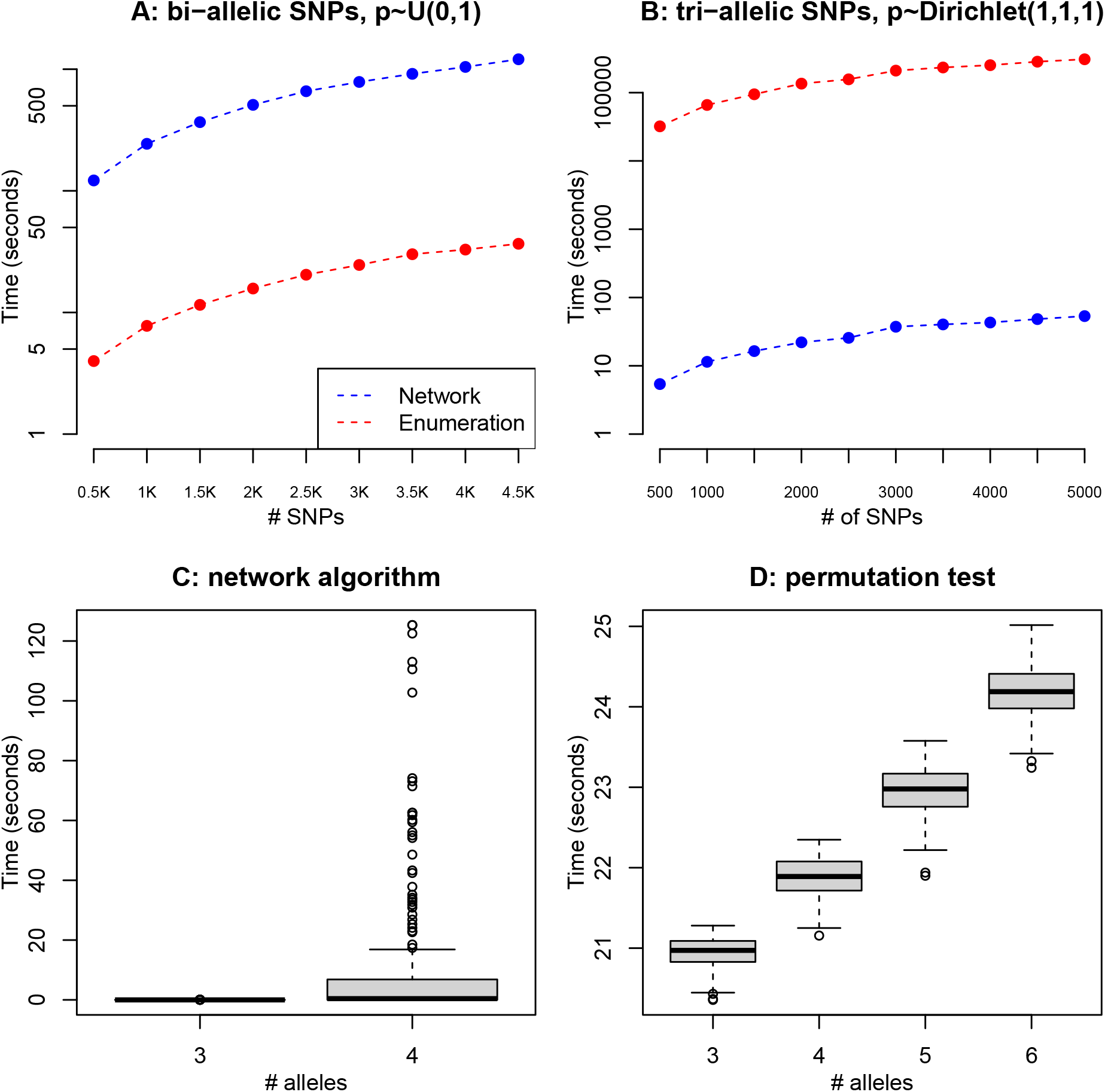
Execution times in seconds for the classical enumeration algorithm, the network algorithm and the permutation test. A: Execution time (in a logarithmic scale) as a function of the number of biallelic SNPs with uniform allele frequencies. B: Execution time (in a logarithmic scale) as a function of the number of triallelic SNPs with Dirichlet(1,1,1) allele frequencies. C: Boxplot of execution times of the network algorithm for 250 SNPs with three and four alleles. D: Execution times of the permutation test for 250 SNPs with three, four, five or six alleles.

These calculations were repeated for simulated triallelic variants, for which the results are shown in Fig. 3B, where we simulated up to 5,000 variants, which is close to the amount of triallelics found on X in the 1,000 genomes project (see Fig. 4A). These figures show that for three-allelic variants the network algorithm is much faster than the enumeration algorithm. We note that it takes the enumeration algorithm 85.5 hours to calculate the maximum of 5,0000 X-triallelics, whereas the network algorithm does this in 53.4 seconds. Execution times also increase linearly with the number of SNPs. For larger numbers of alleles an enumeration algorithm is currently not available. For three through six alleles we compare the network algorithm with the permutation test. We generated 250 multiallelic polymorphisms for a given number of alleles under the assumption of equal allele frequencies in the sexes and Hardy-Weinberg proportions for females, using the Dirichlet distribution with all concentration parameters equal to 1 to simulate the allele frequencies. Figure 3C and 3D shows boxplots of the execution time (in seconds) for the 250 simulated variants as a function of the number of alleles for both the network and permutation tests. The execution time of the permutation tests only experiments a minor increase when the number of alleles increases from three to four. For three and four alleles the network algorithm generally provided the fastest solution. For four alleles, Fig. 3C shows some hard polymorphisms appear for which the network needs more time than a permutation test. On average, the network algorithm is much faster, and outperforms the permutation test with 17,000 draws. Beyond four alleles the permutation test is feasible for all polymorphisms, whereas the computation cost of the network algorithm becomes prohibitive.

**Figure 4:**
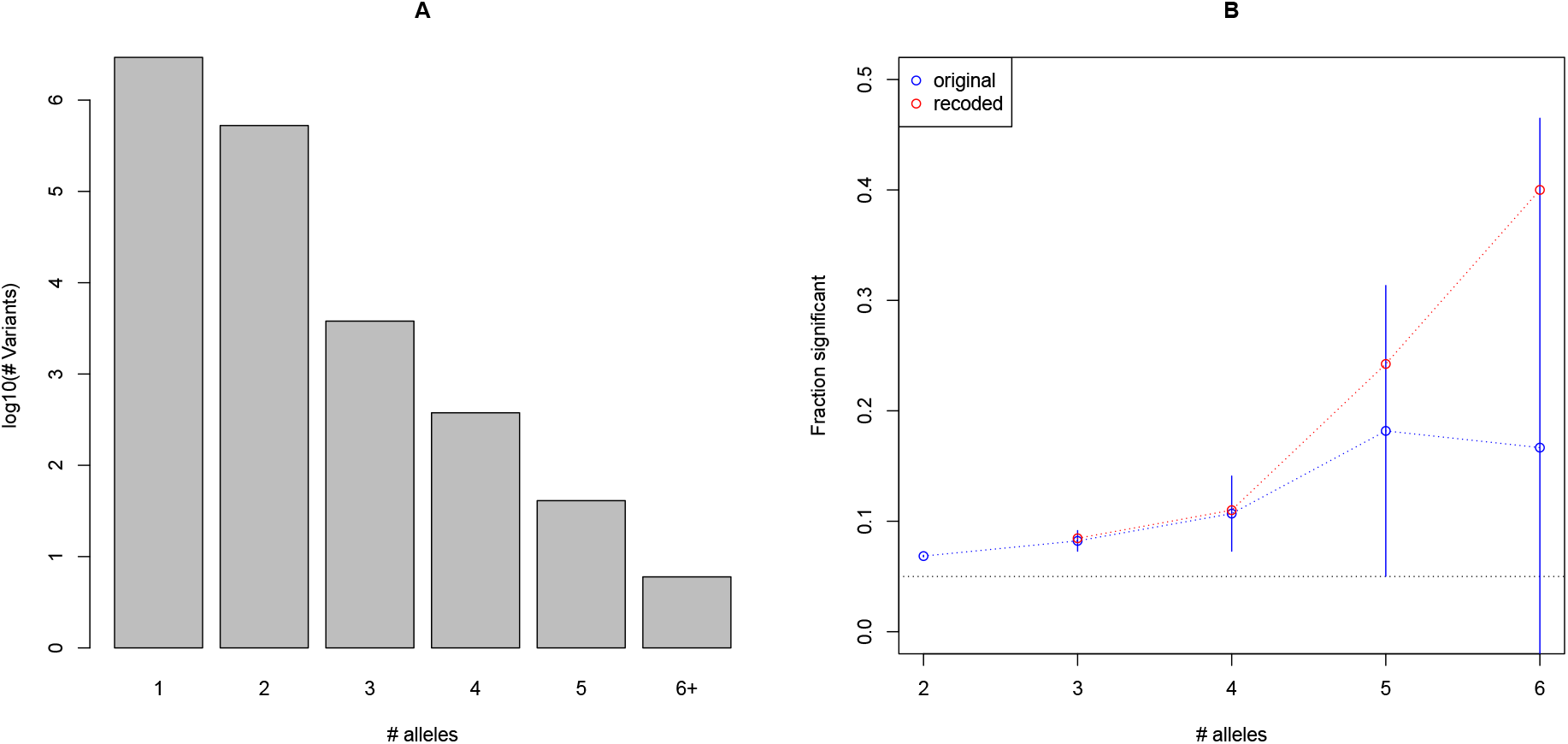
A: Number of variants with a given number of alleles for the TSI population. B: Fraction of significant variants for a given number of alleles. Vertical lines represent 95% confidence intervals for the theoretical fraction. The horizontal dotted reference line represents the significance level *α* = 0.05. Blue open dots represent observed fractions of significant variants. Red open dots present observed fractions of significant variants when the polymorphism is recoded as biallelic.

## 4 Empirical data examples

We present examples of the application of X chromosomal exact tests based on the network algorithm for multiallelic variants taken from the 1,000 genomes project (The 1000 Genomes Project Consortium, 2015) and for a forensic database of X-chromosomal microsatellites (Chen et al., 2018).

### 4.1 TSI sample of the 1,000 genomes project

We consider the analysis of a complete X chromosome of a sample of the TSI population (Tuscany, Italy) of the 1,000 genomes project, using all its multiallelic variants stored in the VCF files of the project, and using the VCFR package (Knaus and Grunwald, 2017) to process the data. This data set consists of 107 individuals, 53 males and 54 females. Variants in the pseudo-autosomal regions (Graves et al., 1998) were excluded from the analysis. Figure 4A shows a barplot with the prevalence of variants with a given number of alleles, and confirms the well-known fact that for a given human population, most variants are monomorphic or biallelic. We calculated the fraction of significant variants (using *α* = 0.05) for each given number of alleles, which reveals that for multiallelic variants more evidence for disequilibrium is found, as shown in Fig. 4B.

Using the enumeration algorithm for all biallelic X-chromosomal variants, the network algorithm for all variants with three through five alleles, and the permutation test to analyse variants with six or more alleles, it took about 10 minutes to analyse all polymorphisms of the TSI sample (*n* = 107); this could be reduced if a few hard three through five allelic variants would be resolved by using the permutation test, at the expense of less precision. We illustrate the observed faster computation of X-chromosomal exact test results with some triallelic polymorphisms. Table 1 shows genotype counts and execution times for six different SNPs. Enumeration and network algorithm produce the same p-value, and the permutation p-value is close to these p-values. For 17,000 permutations, the permutation test takes about one minute to complete. The enumeration algorithm is faster than the permutation test for those variants that have a dominant major allele. For variants rs200225892 and rs11439044 alternate alleles have substantial counts and in these cases the permutation test is faster than full enumeration. In all cases the network algorithm outperforms the permutation and enumeration tests. The network algorithm also requests more computation time for the two variants with larger alternate allele frequencies.

**Table 1:**
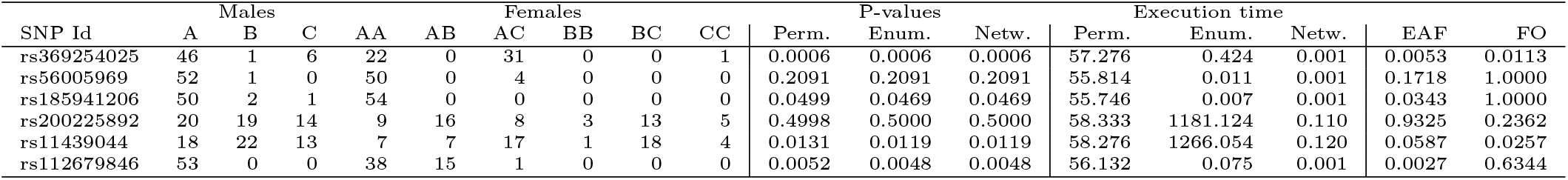
Genotype counts, p-values and execution times (in seconds) for permutation, enumeration and network algorithm of six SNPs of the TSI sample of the 1,000 genomes project. Exact p-values for a test of equality of allele frequencies (EAF) and HWP in females only (FO) are also reported.

Interpreting the genotype patterns, one sees that for rs369254025 HWE is rejected because of different allele frequencies for the sexes and excess heterozygosity for females; for rs56005969 no significant deviations are found; for rs185941206 HWE is rejected because females are monomorphic, whereas males carry all three alleles; For rs200225892 all three alleles are common and no significant deviations are found; for rs11439044 females are out of HW proportions; Finally, for rs112679846 males are monomorphic, but females have a large number of alternate alleles. Notice that disequilibrium would have gone unnoticed for variants rs185941206 and rs112679846 if equilibrium would have been tested in females only.

### 4.2 X-chromosomal STRs of Han Chinese

We use a forensic database of 19 X-STRs of 206 unrelated Han Chinese individuals form Guizhou (104 females and 102 males) described by Chen et al. (2018). Table 2 gives the p-values of permutation tests and network algorithm exact tests for HWE along with the execution time. The X-chromosomal exact tests for all individuals was used, as well as an autosomal test that uses the females only. On average, the X-chromosomal permutation test with 17,000 draws takes about 52 seconds to complete. We observe good agreement between the p-values obtained by the permutation test and by the the network algorithm. The X-chromosomal network algorithm is seen to be much faster for a four-allele STR, slower for a five-allele STR, and not feasible for the remaining STRs which have 7+ alleles.

**Table 2:** Test results and execution times for X-chromosomal STRs. STR identifier, number of STR alleles, permutation test p-value and execution time, network based exact test and execution time, and the same test results based on an autosomal test for the females only, for 19 X-STRs. Dashes (-) represent results not available for requiring too much computation time. Execution times are expressed in seconds (s), minutes (m) or hours (h) as convenient.

| STR | # alleles | All individuals |  |  |  | Females only |  |  |  |
| --- | --- | --- | --- | --- | --- | --- | --- | --- | --- |
|  |  | Permutation |  | Network |  | Permutation |  | Network |  |
|  |  | p-value | time | p-value | time (s) | p-value | time (s) | p-value | time (s) |
| 1 DXS8378 | 5 | 0.0355 | 42s | 0.0345 | 951 | 0.0548 | 40s | 0.0545 | 0.012s |
| 2 DXS7423 | 4 | 0.5171 | 40s | 0.5099 | 0.3 | 0.2678 | 39s | 0.2628 | 0.003s |
| 3 DXS10148 | 17 | 0.4152 | 69s | - | - | 0.3905 | 67s | - | - |
| 4 DXS10159 | 10 | 0.0768 | 49s | - | - | 0.0846 | 48s | - | - |
| 5 DXS10134 | 16 | 0.3472 | 64s | - | - | 0.6156 | 62s | - | - |
| 6 DXS7424 | 10 | 0.9619 | 48s | - | - | 0.8616 | 47s | 0.8573 | 37.1h |
| 7 DXS10164 | 9 | 0.5226 | 46s | - | - | 0.3258 | 45s | 0.3238 | 77.0s |
| 8 DXS10162 | 10 | 0.6986 | 48s | - | - | 0.3981 | 47s | 0.4025 | 6.9h |
| 9 DXS7132 | 8 | 0.7882 | 45s | - | - | 0.5960 | 45s | - | - |
| 10 DXS10079 | 10 | 0.6003 | 49s | - | - | 0.8403 | 49s | - | - |
| 11 DXS6789 | 9 | 0.4776 | 47s | - | - | 0.7255 | 47s | - | - |
| 12 DXS101 | 12 | 0.1845 | 54s | - | - | 0.1572 | 53s | - | - |
| 13 DXS10103 | 7 | 0.8630 | 44s | - | - | 0.8957 | 44s | - | - |
| 14 DXS10101 | 19 | 0.0456 | 75s | - | - | 0.0082 | 73s | - | - |
| 15 HPRTB | 8 | 0.7551 | 45s | - | - | 0.3819 | 44s | 0.3800 | 8.1m |
| 16 DXS6809 | 10 | 0.1722 | 49s | - | - | 0.1194 | 49s | - | - |
| 17 DXS10075 | 9 | 0.4762 | 46s | - | - | 0.3136 | 46s | 0.3109 | 72.2m |
| 18 DXS10074 | 11 | 0.6746 | 50s | - | - | 0.1791 | 49s | - | - |
| 19 DXS10135 | 21 | 0.5691 | 83s | - | - | 0.1308 | 82s | - | - |

The permutation test is, as expected, slightly faster for tests that use females only because of a smaller number of alleles. Application of the network algorithm to the females only leads to spectacular savings in computation time for the two STRs with four and five alleles; for seven or more alleles the permutation test outperforms the network algorithm. Two STRs, DXS8378 and DXS10101 appear as significant at the 5% level; DX8378 for having different allele frequencies in the sexes (*p* = 0.022), and DXS10101 for having females out of Hardy-Weinberg proportions (*p* = 0.008).

## 5 Discussion

We have developed a network algorithm for the X-chromosomal exact test for Hardy-Weinberg equilibrium with multiple alleles. X-chromosomal exact tests were hitherto only feasible for two or three alleles by using an classical full enumeration algorithm. For analysing variants with more alleles a permutation test was required. The network algorithm proposed in this paper extends the feasibility of the X-chromosomal exact test. It is now possible to obtain exact p-values for triallelic X chromosomal variants within fractions of a second (see Table 1). In general, for variants with over four alleles, the computational cost of the network algorithm is still prohibitive, and one still needs to resort to a permutation test or Markov chain approach to resolve these cases. X chromosomal STRs with over four alleles are common, and it remains a challenge to further improve algorithms for obtaining exact instead of approximate p-values in this setting. In forensics, where STRs with many alleles are widely used, a permutation test or Markov chain algorithm thus remain the best general purpose methods that will serve for all STRs. The network algorithm may be more interesting for the analysis of indels (Mills et al., 2006), which have in general a much smaller number of alleles.

In exact testing for HWE with multiallelic genetic variants *tied outcomes* can easily arise. A pair of tied outcomes refers to two different genotype arrays that have theoretically exactly the same probability under the equilibrium null distribution. Tied outcomes can be problematic if they involve the observed sample. If a genotyping array has the same probability as the observed sample, its probability should be included in the calculation of the p-value. Due to finite precision in the comparison of floating point numbers on a computer, a theoretically tied outcome may not be recognised as having the same probability as the observed sample, and may eventually not be counted towards the p-value. A good computational strategy for comparing probabilities and deciding upon equality of floating point numbers is therefore crucial for a correct implementation of exact test procedures. A permutation test will not resolve the problem of ties, because it also relies on the comparison of the probability of the observed sample with those of other, possibly tied outcomes generated under the null distribution.

Figure 4B shows more evidence against HWE for multiallelic variants. At first sight this may suggest multiallelic variants are more prone to genotyping error. This is, however, hard to tell because the statistical power of tests for disequilibrium depends on the distribution of the allele frequencies, and tests have less power at low MAF (Wigginton et al., 2005; Graffelman and Moreno, 2013). On the other hand, if all multiallelic variants are recoded as biallelic (A the most common allele, B any other allele) then the fraction of significant variants remains increasing with the number of alleles, finally indicating that there is apparently more disequilibrium in multiallelic variants.

Graffelman and Moreno (2013) proposed the use of the mid p-value in exact tests for HWE, in a biallelic setting, for having a rejection rate that is closer to the nominal significance level. In the current multiallelic setting, the mid p-value can easily be obtained by subtracting half the probability of the observed sample, using Eqs. 1 and 4 for the autosomal and X chromosomal case respectively, from the standard exact p-value obtained by the network or permutation algorithm.

The comparison of execution times of different algorithms reflects the performance of current state-of-the-art functions in the R environment. The comparison is facilitated by the fact that all execution time measurements were made inside the R environment on the same linux cluster. However, observed differences are not only due to the algorithm being used, but inevitably also to the coding of the algorithms. It is well known that loops are slower in R than in C, C++ or Fortran, and consequently many R programs can be speeded up by recoding parts in one of these programming languages. In Fig. 3B on triallelic variants we used an enumeration algorithm which was fully written in R, whereas large part of the network algorithm was written in C. Therefore, the better execution time of the network algorithm can at least in part be ascribed to its coding in C. If the enumeration algorithm had been coded in C, a less striking difference between the two algorithms would have been observed.

In summary, we have made progress obtaining a great computational improvement for exact HWE testing at three and four allelic X chromosomal variants. The advantage of exact tests is that they do not rely on approximation. It remains a challenge to further improve algorithms and coding for the exact testing of variants with more alleles.

## 6 Software

The network algorithm for the X-chromosomal exact test with multiple alleles is implemented in function `HWNetwork` of version 1.6.7 of the R-package `HardyWeinberg` (Graffelman, 2015). We adapted C code for an autosomal network algorithm from Engels’ `HWxtest` package available at https://github.com/wrengels/HWxtest. We imported C functions into R through the `Rcpp` package (Eddelbuettel and Francois, 2011).

## Acknowledgements

This work was supported by grant RTI2018-095518-B-C22 (MCIU/AEI/FEDER) of the Spanish Ministry of Science, Innovation and Universities and the European Regional Development Fund, and by grant R01 GM075091 from the United States National Institutes of Health.

## Author contributions

JG conceived the analysis and the article. LO wrote computer programs in C. Both authors performed data analysis, and contributed to the writing of the article.

## Data accessibility

The TSI genotype data is available at the www.internationalgenome.org. The example polymorphisms in Table 1 have been included as a data object `TSI_X_TriAllelics` in R package `HardyWeinberg`. The X-chromosomal STR Guizhou Han data used in Section 4.2 is available as supporting information of the article by Chen (2018).

## Notes

### Competing Interest Statement

The authors have declared no competing interest.

